# Using dynamic ultrasound to assess Achilles tendon mechanics during running: the effect on running pattern and muscle-tendon junction tracking

**DOI:** 10.1101/2024.08.23.609380

**Authors:** Wouter Schallig, Ytjanda Sloot, Milou M. van der Schaaf, Sicco A. Bus

## Abstract

Achilles tendon strain can be quantified using dynamic ultrasound, but its use in running is limited. Minimal effects on running pattern and acceptable test-retest reliability of muscle-tendon junction (MTJ) tracking are prerequisites for ultrasound use during running. We aimed to assess (i) the effect of wearing an ultrasound transducer on running pattern and (ii) the test-retest reliability of MTJ tracking during running. Sixteen long-distance runners (nine injury-free, seven with Achilles tendinopathy) ran at different speeds on an instrumented treadmill with a 10-camera system tracking skin-mounted retroreflective markers, first without and then with an ultrasound transducer attached to the lower leg to track the MTJ of the gastrocnemius medialis. Spatiotemporal parameters, joint kinematics and kinetics were compared between conditions using mixed ANOVAs and paired t-tests. MTJ tracking was performed manually twice by three raters in ten participants. Variability and standard error of measurement (SEM) quantified the inter- and intra-tester test-retest reliability. The running pattern was not affected by wearing the ultrasound transducer, except for significantly less knee flexion during midstance (1.6°) and midswing (2.9°) found when wearing the transducer. Inter-rater and intra-rater SEMs for MTJ tracking to assess the tendon strain (0.43%, and 0.56%, respectively) were about four times as low as between-group differences presented in literature. The minimal effects found on the running pattern and acceptable test-retest reliability indicates that dynamic ultrasound during running can be appropriately used to study Achilles tendon mechanics and thereby help improve our understanding of Achilles tendon behavior during running, injury development and recovery.

## INTRODUCTION

The Achilles tendon is of major importance during running and its tendinopathy has the highest incidence proportion (i.e. 10.3%) of all running-related injuries (Kakouris, Yener and Fong, 2021). The mechanical properties and behavior of the Achilles tendon, like stress, elongation or strain, are associated with tendon failure and tendinopathy (Arya and Kulig, 2010; Finni and Vanwanseele, 2023; Wren et al., 2003). Tendon strain can be quantified during movement using B-mode ultrasound (i.e. dynamic ultrasound) by tracking the muscle-tendon junction (MTJ) between the Achilles tendon and either of the triceps surae muscles. It can therefore have value in understanding and monitoring Achilles tendon injury mechanisms.

Dynamic ultrasound has been applied to walking and other functional tasks in non-injured and injured individuals as well as in pathological conditions (Arya and Kulig, 2010; Cenni et al., 2020; Child et al., 2010; Kalsi, Fry and Shortland, 2016), but its application during running is limited (Farris, Trewartha and McGuigan, 2012; Kharazi et al., 2021). This might be due to methodological challenges. First, wearing an ultrasound transducer might affect the running pattern. It can be perceived as uncomfortable and it is shown to slightly (<5°) affect gait kinematics during normal walking on a treadmill (Mooijekind et al., 2023) and over ground (Cenni et al., 2024). The higher velocity during running might amplify these differences. Second, MTJ tracking can be challenging during running. Higher impact forces increase the soft tissue artifacts and movement due to inertia effects of the marker clusters (Leardini et al., 2005; Wang et al., 2023). Additionally, the higher velocity during running compared to walking will result in faster tendon stretch, reducing MTJ tracking accuracy (Cenni et al., 2020).

Minimal effects on running kinematics and kinetics and acceptable reliability of MTJ tracking are prerequisites for ultrasound use during running to assess Achilles tendon mechanics. Therefore, we aimed to assess (i) the effect of wearing an ultrasound transducer attached to the lower leg on the running pattern and (ii) the inter- and intra-rater test-retest reliability of MTJ tracking during running of injury-free and injured runners. Both groups are included to assess potential group-related differences and ensure the results’ generalizability.

## METHODS

Sixteen participants (six females, mean±SD age:35.4±13.1 years, height:1.81±0.12 m, weight:73.2±11.4 kg, weekly training distance:40.5±18.1 km) of whom nine were injury-free and seven were diagnosed with Achilles tendinopathy (AT) (de Vos et al., 2021) were included. All were experienced long distance runners (i.e. ≥20 km/week for ≥2 years prior to the measurement). Requirement for ethical review of the study was waived by the medical ethics committee of Amsterdam UMC (registration number:#2022.0458). Participants signed written informed consent prior to inclusion.

### Data collection

Participants performed a running protocol on an instrumented treadmill (GRAIL, Motek ForceLink BV, The Netherlands), while wearing their own running shoes. Marker data were collected according to the human body model (Flux et al., 2020; van den Bogert et al., 2013) with a 10-camera optical system (frequency: 100Hz, Vicon Motion Systems, Oxford, UK). Additionally, a rigid 4-marker cluster was attached to the calcaneal dorsolateral side to track the calcaneus instead of the shoe (Figure 1A-B). The position of the Achilles tendon insertion, determined with the subject prone and a neutrally positioned ankle, was expressed in the calcaneal cluster coordinate system by using a pointer with the subject standing (Figure 1A-C). Another rigid 4-marker cluster was attached to the ultrasound transducer (Figure 1A-D) to obtain position and orientation of the ultrasound images in space and thereby the MTJ location (i.e. Achilles tendon origin).

**Figure 1.**
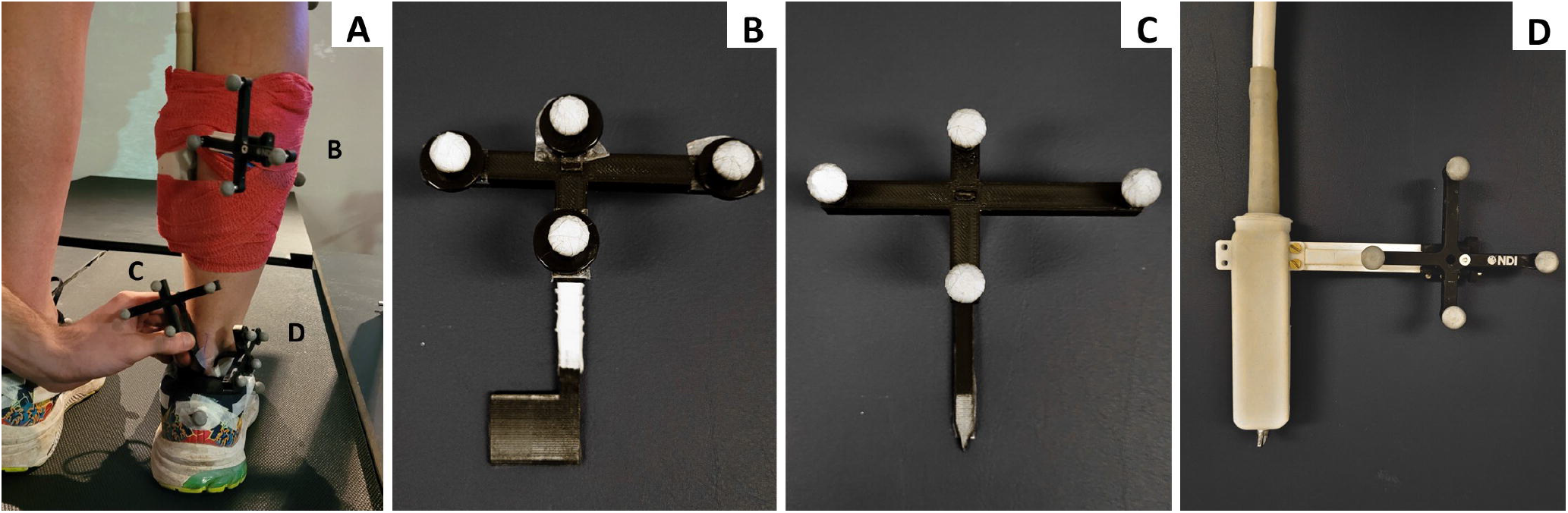
Overview of the marker clusters used during the measurements to be able to quantify the Achilles tendon length during static and dynamic trials (A), with the rigid cluster used to track the calcaneus (B), a pointer with 4 markers attached to indicate the position of the tendon insertion (C) and a rigid cluster attached to the ultrasound transducer with probe holder (D).

During a static standing trial in anatomical position, the segment coordinate systems and Achilles tendon reference length were determined. Next, successive two-minute blocks of running at 10 km/h, 12 km/h, 90% of their own 10 km race pace and, if possible, 14 km/h were performed. After a 10-minute break, the same running bouts were performed with a 59 mm linear ultrasound transducer (LV8-5L60N-2, Telemed, SmartUS, Lithuania, ±110 grams) attached in the longitudinal direction of the lower leg (i.e. injured or randomly selected side for AT and injury-free runners, respectively) at the level of the MTJ of the m. gastrocnemius medialis using self-adhesive tape (Figure 1A). The transducer operated at 31 frames per second for optimal single image quality (depth:50 mm, dynamic range:72 dB, power:-2 dB, gain:58%, line density:standard, frequency:8 MHz). Data were collected for 20 seconds after one minute of running at each speed.

### Data processing: gait analysis

Only the fastest speed trial was processed because it was considered most challenging. Marker and force data were processed with Vicon Nexus (v.2.11, Oxford, UK) and spatiotemporal parameters, joint kinematics and kinetics were computed using the Gait Offline Analysis Tool (GOAT v.4.2, Motek, the Netherlands). Gait events were quantified following Zeni et al. (2008). Per participant, five random strides were selected and averaged. Spatiotemporal parameters, joint kinematics and kinetics were compared between conditions using either regular or 1D statistical parametric mapping ANOVAs to test for the effect of transducer attachment, injury status and their interaction effect, followed by post-hoc t-tests. For the statistically significantly different parts of the gait cycle, root mean square differences were quantified (RMSD). All tests were performed in Matlab (R2017b, Mathworks, USA) with significance level set at α=0.05.

### Data processing: ultrasound

Ultrasound images of the static and dynamic 12 km/h trials were processed in ImageJ (v.1.53, Bethesda, MD), by using the multi-point function the MTJ coordinates in each image were manually selected. For both the static and dynamic trials, a customized Python algorithm transformed the local MTJ coordinates to global coordinates, to determine Achilles tendon length by calculating the distance between insertion and origin (Cenni et al., 2016). Achilles tendon strain (%) was calculated in each ultrasound frame using equation 1 (Farris, Trewartha and McGuigan, 2012), where ATL and ATL_ST_ represent the dynamic and static Achilles tendon length, respectively.

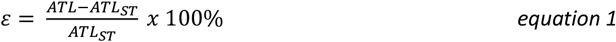

Three independent raters processed the data twice for 10 randomly selected participants. At least 24 hours was between repeated data processing assessments in each participant. Data of the other six participants were used beforehand as practice dataset, to reach consensus in allocating the MTJ. Intra-rater and inter-rater (using the second assessment of each rater) test-retest reliability were determined for strain, and dynamic and static tendon length. Variability and standard error of measurement (SEM) were used as reliability measures. Variability was calculated as the standard deviation at each percent of the stance phase for each pair (intra) or trio (inter) of MTJ measurements. These were averaged over the stance phase of 10 consecutive strides to obtain one variability value (σ) for each participant. Next, SEM was calculated as the root mean square average of the 10 participants, leading to inter-tester reliability and intra-tester reliability for each tester separately (Bland and Altman, 1996). Intra-rater reliability for the study was the average of the three testers.

## RESULTS

The trials analyzed for the effect on running pattern were performed at 13.3±1.5 km/h. From the total of 16 participants, two were excluded from analysis due to treadmill malfunction or missing markers. Two more were excluded from kinetic data analysis due to ground reaction force errors.

No significant effect of condition (Table 1.), injury status nor an interaction effect were found for step width, stride length and stride time.

**Table 1.**
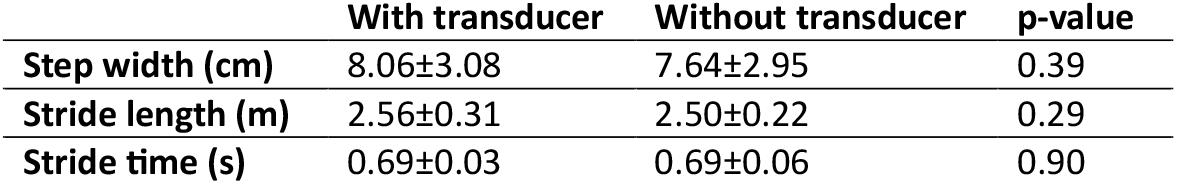
Spatiotemporal parameters for running with and without ultrasound transducer.

Running kinematics and kinetics were mostly not significantly different between conditions (Figure 2-3), nor was there any significant effect of injury status or interaction effect. The knee was significantly less flexed during midstance (RMSD=1.6°) and midswing (RMSD=2.9°) with the ultrasound transducer attached. Additionally, for a brief period in the swing phase (71.3-74.6%) the ankle was less supinated (RMSD=2.3°) with the ultrasound attached. Only during brief phases (i.e. <4%, mainly in swing) the kinetics were significantly different between conditions.

**Figure 2.**
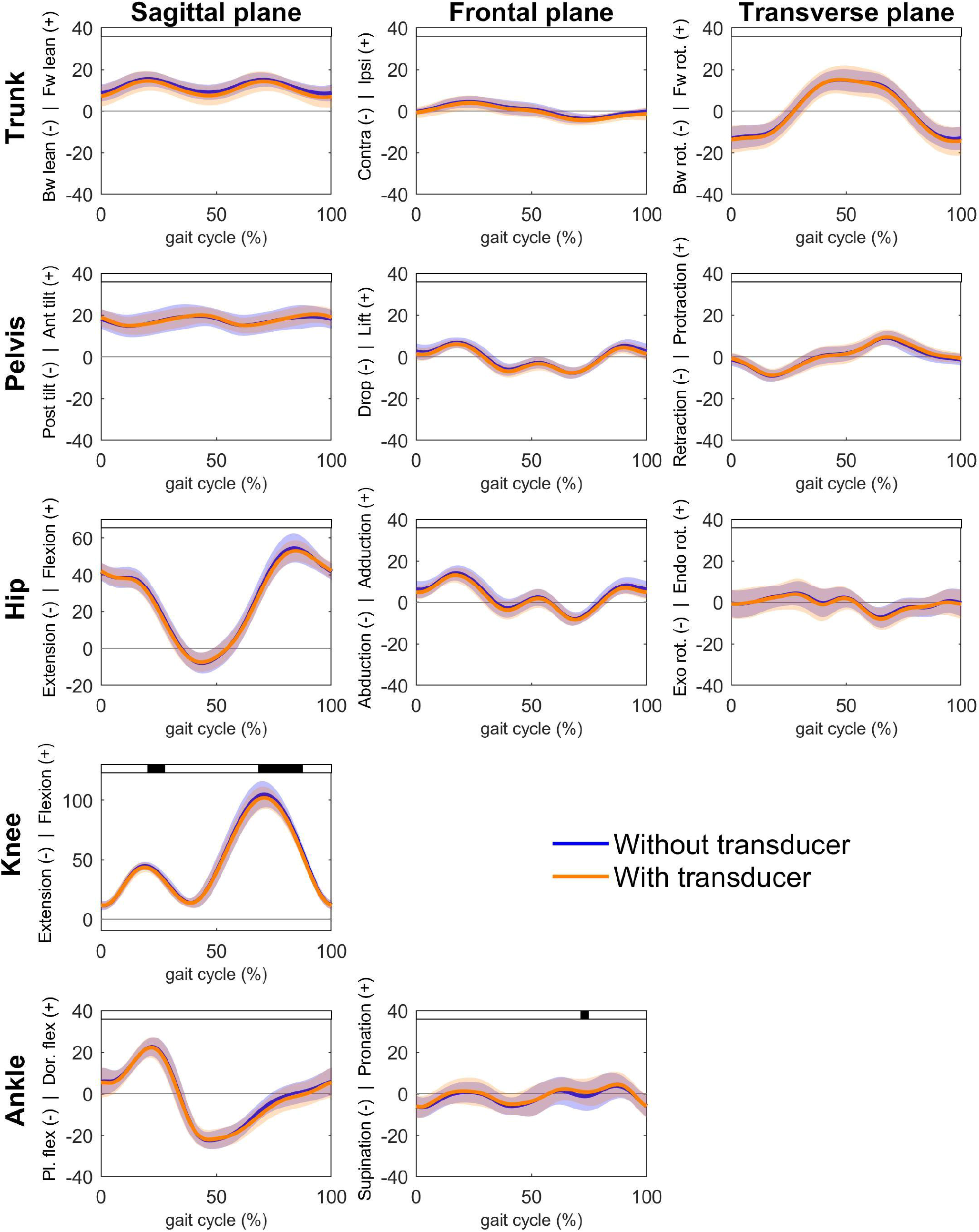
Kinematic comparison between the conditions without (blue) and with (orange) an ultrasound transducer attached to the lower leg. Black bars at the top depict statistically significant differences between running conditions. *Abbreviations: Bw: backward, Fw: forward, Post*.: *posterior, Ant: anterior, Pl. flex: plantar flexion, Dor. flex: dorsiflexion, Exo rot: exorotation, Endo rot: endorotation*.

**Figure 3.**
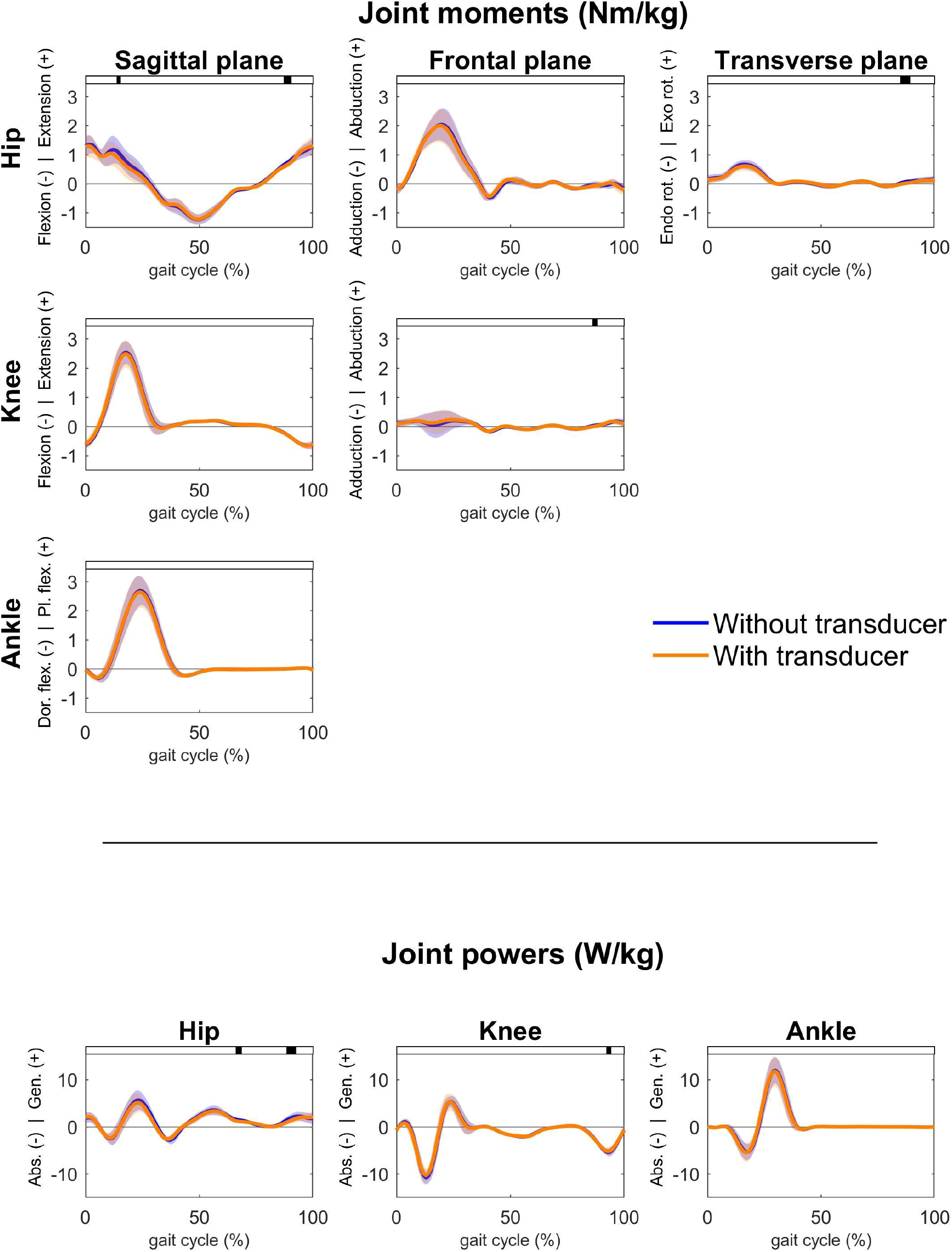
Kinetic comparison between the conditions without (blue) and with (orange) an ultrasound transducer attached to the lower leg. Black bars at the top depict statistically significant differences between running conditions. *Abbreviations: Pl. flex: plantar flexion, Dor. flex: dorsiflexion, Exo rot: exorotation, Endo rot: endorotation, Abs: Absorption, Gen: Generation*.

Intra-rater test-retest reliability of the MTJ tracking was higher than inter-rater reliability for the tendon strain, while vice versa for the static and dynamic tendon length (Table 2). Static tendon length showed higher errors than dynamic tendon length for both types of reliability. SEM for the three individual raters were 0.33, 0.58 and 0.77% for the strain, 0.86, 0.42 and 0.34 mm for the dynamic length and 0.35, 0.89 and 1.13 mm for the static length. Appendix A provides the average time series for additional insight.

**Table 2.** Reliability measures of the muscle-tendon junction tracking for the tendon strain, the dynamic tendon length during running and the static tendon length during standing.

|  |  | Strain [%] | Dynamic length [mm] | Static length [mm] |
| --- | --- | --- | --- | --- |
| **Inter-rater** | $\sigma$ | 0.36 | 0.61 | 0.58 |
|  | SEM | 0.43 | 0.69 | 1.03 |
| **Intra-rater** | $\sigma$ | 0.34 | 0.42 | 0.42 |
|  | SEM | 0.56 | 0.54 | 0.79 |

## DISCUSSION

Our main findings were that the running pattern of injury-free and injured runners was mostly not affected by wearing an ultrasound transducer on the lower leg, except for less knee flexion during midstance and midswing. Furthermore, inter-rater and intra-rater test-retest SEM values in tendon strain were <0.6%, in static length <1.1 mm and dynamic length <0.7 mm.

Running with an ultrasound transducer attached to the lower leg showed similar results as found during normal walking, for which reduced knee flexion during midswing was also shown (Cenni et al., 2024; Mooijekind et al., 2023). probably due to the ultrasound cable that closely passes the knee joint. These swing phase kinematics might not be most relevant in the context of assessing Achilles tendon mechanics, where the stance phase with its higher tendon loads is more relevant. We also found less knee flexion during midstance, potentially resulting in a lengthened muscle-tendon unit compared to running without transducer. Nevertheless, differences were <5°, generally considered the limit for clinically acceptable errors in gait analysis (McGinley et al., 2009). In the kinetic data, only brief and small differences during swing were found between conditions. These differences were possibly due to the additional weight of the transducer (±110 grams), but considered clinically irrelevant because they were minor and present during swing phase.

This study is one of the first to report the reliability of manual digitization of the MTJ in ultrasound images during running, demonstrating similar values for dynamic tendon length as one other study which reported errors <0.5 mm in x or y direction (Krikelis, Pain and Furlong, 2021). Our variability and SEM values for strain were smaller for inter-rater compared to intra-rater reliability, while the SEM for dynamic and static length were both larger for the inter-rater reliability. A possible explanation is that for each rater the inconsistency in MTJ tracking for static and dynamic length is in the same direction, which are cancelled out when calculating the strain. Strain reliability was mainly affected by static trial differences. Important to note is that with two testers one extreme variability value was present, for both in the same participant, which largely affected the average intra-rater reliability for the static trial for these two raters. Excluding that single participant would reduce their intra-rater SEM from 0.89 and 1.13 to 0.23 and 0.29 mm, respectively. This highlights the importance of proper MTJ quantification in the static trial. Moving the ankle through plantar-dorsiflexion motion at the beginning or end of the static trial might help to get a better indication of the MTJ location.

Test-retest reliability values should be placed into perspective by comparing them to actual strain and tendon length values during running. Achilles tendon peak strain values are reported to be 4% during walking compared to 4.9% during running (Kharazi et al., 2021), and strain differences between injured and non-injured runners of 1.8% have been found for maximal isometric plantar flexion (Child et al., 2010). MTJ displacement during running is about 20 mm at 12.6 km/h (Kharazi et al., 2021), which is similar to the 12.0 km/h used in the current study. The SEM for strain we found are about 3-4 times as low as between-group differences, and for tendon length 18x as small as displacements presented in literature. Hence, assessments seem reliable and acceptable for their purpose.

Several factors should be considered regarding generalizability of our results. First, these results are applicable to gastrocnemius medialis MTJ tracking, using dynamic ultrasound for other triceps surae muscles (Adam et al., 2023) requires different probe locations and potentially more challenging MTJ tracking. Second, the ultrasound transducer operated at a lower frequency (31 fps) than used in other studies to improve single image quality, potentially affecting tracking reliability over time, because only about 20 images are analyzed per stride. Hence sufficient gait cycles have to be included in the analysis to obtain reliable strain waveforms. Nevertheless, we have found similar Achilles tendon length and strain waveforms (Appendix A) as other literature (Farris et al., 2012; Kharazi et al., 2021). Third, we tracked the MTJ manually, while (semi-)automatic tracking software is available. When the larger errors from more automated tracking (Cenni et al., 2020) are reduced, it is preferred for future applications to reduce processing time. Fourth, we focused solely on the MTJ tracking reliability, but transducer placement can also introduce reliability errors due to e.g. probe tilt (Van Hooren, Teratsias and Hodson-Tole, 2020) which might add to the measurement error. Fourth, since we did not find an effect of injury status on the effect of the transducer on the gait pattern, nor on the reliability analysis, the results are applicable to the injured and non-injured runners.

In conclusion, the running pattern is mostly not affected by wearing an ultrasound transducer on the lower leg and the inter- and intra-rater test-retest reliability of tracking the MTJ of the medial gastrocnemius is within acceptable ranges. Therefore, we advocate the use of dynamic ultrasound during running to study Achilles tendon mechanics and thereby help further improve our understanding of Achilles tendon behavior during running, injury development and recovery.

## Acknowledgements

The authors would like to thank Mabel Brands and Sander de Hoog, Masters students Human Movement Sciences, for their help in the data collection.

