## Appendix A for "Using dynamic ultrasound to assess Achilles tendon mechanics during running: the effect on running pattern and muscle-tendon junction tracking"

This paper focused on the intra- and inter-repeatability of the Achilles tendon strain. To gain additional insight in the strain and length data we have plotted the average and standard deviation (shaded area) over participants and testers in the figure below for the tendon strain, dynamic tendon length and static tendon length (dashed line).

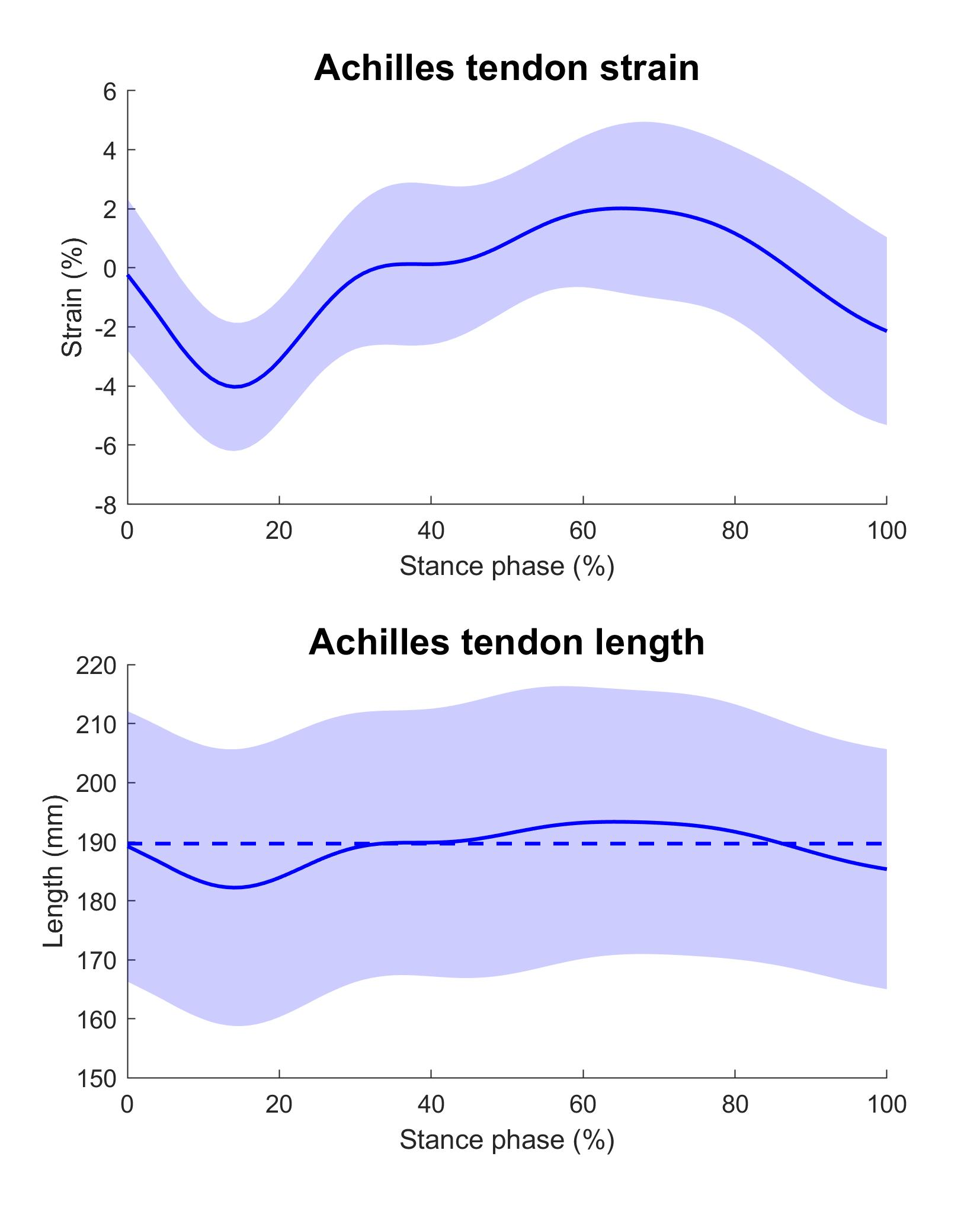
